# Dynamic dissociation of the IFT complex drives ciliary dysfunction during *C. elegans* ageing

**DOI:** 10.64898/2026.04.15.718672

**Authors:** Jiayi Li, Songyue Wang, Shengzhi Xie, Meng Chen, Mengjiao Song, Xiaona Zhang, Ji Tang, Qifu Li, Dong Li, Xiumin Yan, Yidong Shen

## Abstract

Intraflagellar transport (IFT) is essential for cilia, and its dysfunction drives ciliopathies and systemic ageing. However, the *in vivo* dynamics of individual components in the IFT complexes remain obscured, leaving the mechanisms of age-dependent IFT failure largely unknown. Here, we report a dual-color super-resolution imaging strategy to dissect the structural integrity and kinetics of IFT trains in the sensory cilia of young and aged *Caenorhabditis elegans*. We show that IFT complexes are not static entities. Instead, distinct components undergo dynamic dissociation within IFT trains. This intra-complex dissociation causes a remarkable reduction in IFT velocity and is significantly exacerbated in the cilia of aged worms. Mechanistically, we identify the conserved TRiC/CCT chaperonin complex and *daf-19*/RFX, the master transcription factor driving IFT genes, as critical regulators of IFT stability. We demonstrate that their age-dependent downregulation drives the progressive IFT component dissociation. Our findings re-frame the IFT complex as a highly dynamic assembly, uncover a new dimension of IFT regulation, and identify the progressive uncoupling of IFT components as a key driver of ciliary dysfunction during ageing.

## Introduction

Cilia are conserved, microtubule-based organelles that project from the surface of most eukaryotic cells, critical for cell motility, fluid flow, and signal transduction. Dysfunction in cilia leads to a spectrum of disorders collectively termed ciliopathies, including polycystic kidney disease, retinal degeneration, neurodevelopmental defects, situs abnormalities, obesity, and infertility (Goetz & Anderson, 2010; Gopalakrishnan *et al*, 2023; Green & Mykytyn, 2010; Tobin & Beales, 2009). Emerging evidence further links cilia to ageing and age-associated pathologies (Oya *et al*, 2024; Silva & Cavadas, 2023; Volos *et al*, 2025).

Most ciliary proteins are synthesized in the cytoplasm, transported into cilia and recycled back into the cell body by intraflagellar transport (IFT). IFT is a bidirectional trafficking system driven anterogradely by kinesin-2 motors and retrogradely by dynein-2. IFT trains comprise two multisubunit complexes, IFT-A (6 subunits) that is required for retrograde transport and IFT-B (16 subunits) is responsible for the anterograde IFT. The BBSome (BBS1, 2, 4, 5, 7, 8, 9, 18) couples membrane cargos into IFT and mediates their retrieval (Cole & Snell, 2009; Lacey *et al*, 2023; Lacey & Pigino, 2024; Mitra *et al*, 2024; Nakayama & Katoh, 2018; Ou *et al*, 2005; Prevo *et al*, 2017). Consistent with these roles, mutations in IFT subunits, motors, or the BBSome perturb ciliary trafficking and architecture, resulting in ciliopathies (Mul *et al*, 2022; Pigino, 2021; Rosenbaum & Witman, 2002).

Live imaging has been central to defining IFT dynamics since its initial application in *Chlamydomonas* by the Rosenbaum lab using differential interference contrast microscopy, and across *Chlamydomonas*, *C. elegans* and mammalian cilia it has quantified velocities, frequencies, and dwell times of IFT trains (Kozminski *et al*, 1993). Super-resolution and correlative imaging approaches resolve parallel transport “lanes” along axonemal microtubules (MTs) in *Chlamydomonas* to avoid collisions between anterograde trains on B-tubules and retrograde trains on A-tubules (Stepanek & Pigino, 2016). Advanced live imaging techniques, including Hessian structured illumination microscopy and grazing incidence structured illumination microscopy (GI-SIM) (Huang *et al*, 2018; Qiao *et al*, 2023; Xie *et al*, 2020), further uncover complex behaviors such as pausing, returns and sidestepping of IFT. Despite these advances, due to the inadequate spatial and temporal resolution, most live-imaging IFT studies rely on single-color reporters, limiting direct observation of how distinct IFT subcomplexes and motors co-move, remodel and load/unload cargo in real time. Moreover, IFT dynamics have been largely characterized in young animals, leaving their modulation under physiological stress such as ageing poorly understood.

We previously found in *C. elegans* that ciliary length and axonemal organization remain grossly intact with age, whereas the frequency and velocity of IFT significantly decline due to the age-dependent reduction of IFT components expression (Zhang *et al*, 2021a). We reasoned that the dysregulation of IFT components in aged worms could disrupt their ratio in IFT complex, promoting in-transit detachment or imperfect coupling among IFT subcomplexes, compromising train integrity and ultimately ciliary sensory function. In this study, we construct a dual-color IFT system to visualize two subunits from IFT-A, IFT-B, motors and the BBSome simultaneously in *C. elegans* using GI-SIM (Cole & Snell, 2009; Hao & Scholey, 2009; Sanders *et al*, 2015) (Fig. 1A). We show that IFT train is not simple base-to-tip-to-base conveyor but exhibits rich dynamics including split, fusion, turnaround, and pause. More strikingly, we find that different train parts occasionally dissociate from each other, reducing IFT velocity. This dissociation increases with age due to the downregulation of TRiC/CCT chaperonin complex, impairing IFT in aged worms.

**Figure 1.**
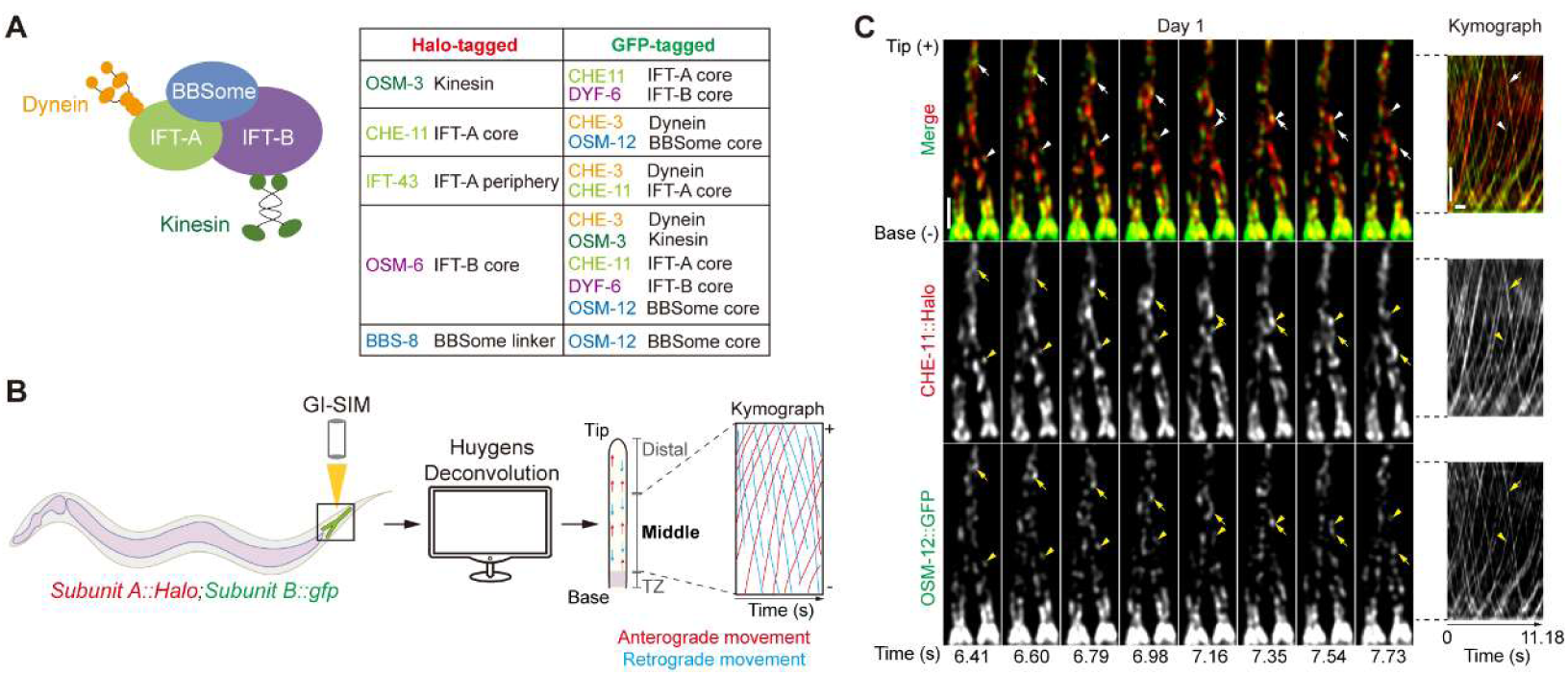
Simultaneous tracking of two IFT components in *C. elegans* cilia. **(A)** The 12 combinations of dual-labeled IFT subunits in this study. **(B)** The workflow of data acquisition and analysis to live track dual-labeled IFT subunits. **(C)** Representative time lapse images and corresponding kymographs of a pair of IFT subunits (CHE-11::Halo and OSM-12::GFP in red and green fluorescence, respectively) in our live imaging assay of a young adult worm at day 1 of adulthood. Arrows and arrowheads denote retrograde and anterograde IFT complexes, respectively. Scale bars: 1 μm in time lapse images. In kymographs, the horizontal scale bars: 1 s, the vertical scale bars: 1 μm. Please also see Video EV1.

## Results

### Simultaneous tracking of distinct IFT components in sensory cilia by GI-SIM

To resolve the real-time dynamics of IFT complexes in living *C. elegans*, we tagged key components of the IFT-A and IFT-B subcomplexes, the BBSome, and the dynein and kinesin motors, which were shown to exhibit age-related declines (Lacey *et al*., 2023; Meleppattu *et al*, 2022; Nakayama & Katoh, 2018; Singh *et al*, 2020; Zhang *et al*., 2021a), with either HaloTag or GFP. We then systematically generated 12 dual-labeled combinations to probe the coordinated movement of components within and between different subcomplexes of the IFT machinery. These combinations included pairings of: (a) IFT-A core and peripheral components; (b) IFT-A core and IFT-B core components; (c) two distinct IFT-B core components; (d) IFT-A/B core components and kinesin; (e) IFT-A/B components and dynein; (f) IFT-A/B core components and a BBSome core component; and (g) a BBSome core component and a BBSome linker protein(Nakayama & Katoh, 2018) (Fig. 1A).

For imaging, the IFT components with HaloTag were labeled in red fluorescence using JF549 ligand. Fluorescent signals from GFP and HaloTag-labeled IFT components were acquired sequentially by GI-SIM at 200 nm spatial resolution for 11.2 sec. 100 frames were captured using an 18 ms acquisition time per channel and an inter-channel switching delay of 1.34 ms. Because the speed of IFT is within 2 μm/s in worm cilia (Zhang *et al*., 2021a), the high spatial-temporal resolution enabled us to live-track a pair of IFT components simultaneously. The raw data were further processed using interleaved reconstruction and Huygens deconvolution to generate high-resolution movies at 27 Hz frame rate and corresponding kymographs of IFT particle trajectories (Fig. 1B,C and Video EV1). Our analysis focused on the 5 μm-long middle segment of the cilia, a region containing microtubule doublets, to ensure unambiguous tracking of individual particles, avoiding the high particle density at the base and complex trajectories at the distal tip.

The velocity and frequency of dual-labeled IFT particles are similar to previous observations using single-labelled imageing (Fig. EV1A,B) (Zhang *et al*., 2021a), indicating that dual labeling does not affect IFT. Moreover, 9 of the 12 combinations showed a decrease in velocity in aged worms, whereas all examined combinations exhibited a reduction in IFT frequency in aged worms (Fig. EV1A,B). These findings are consistent with our previous observations of 4 single-labeled IFT components (Zhang *et al*., 2021a), showing that the age-related decline in transport efficiency is a general feature of the entire IFT machinery.

### A systematic analysis of IFT train behavior in *C. elegans* sensory cilia

Previous reports have shown a series of IFT train behaviors, including ‘split’ of one train into several smaller ones, trains ‘fusion’, intra-ciliary ‘turnaround’, and ‘pause’ of train movement (Bertiaux *et al*, 2018; Buisson *et al*, 2013; Dentler, 2005; Mijalkovic *et al*, 2017; Qiao *et al*., 2023; Stepanek & Pigino, 2016; Sun *et al*, 2025; Zhang *et al*, 2021b). We systematically examined these behaviors in young and aged worms. Because of the relative weak signal of the retrograde trains (Bertiaux *et al*., 2018; Chien *et al*, 2017), our analysis focused on the anterograde IFT.

All four behaviors were observed (Fig. 2 and Videos EV2-5). The incidences of ‘split’ and ‘fusion’ events varied considerably among different subunit pairs but did not show a consistent change between young and aged worms (Fig. 2A-D, Fig. EV2A,B and Videos EV2,3). Unlike ‘split’ and ‘fusion’, the incidence of ‘turnaround’ and ‘pause’ showed a significant increase in aged worms (Fig. 2E-H, Fig. EV2C,D and Videos EV4,5). Moreover, the duration of pause generally increased with age (Fig. EV2D).

**Figure 2.**
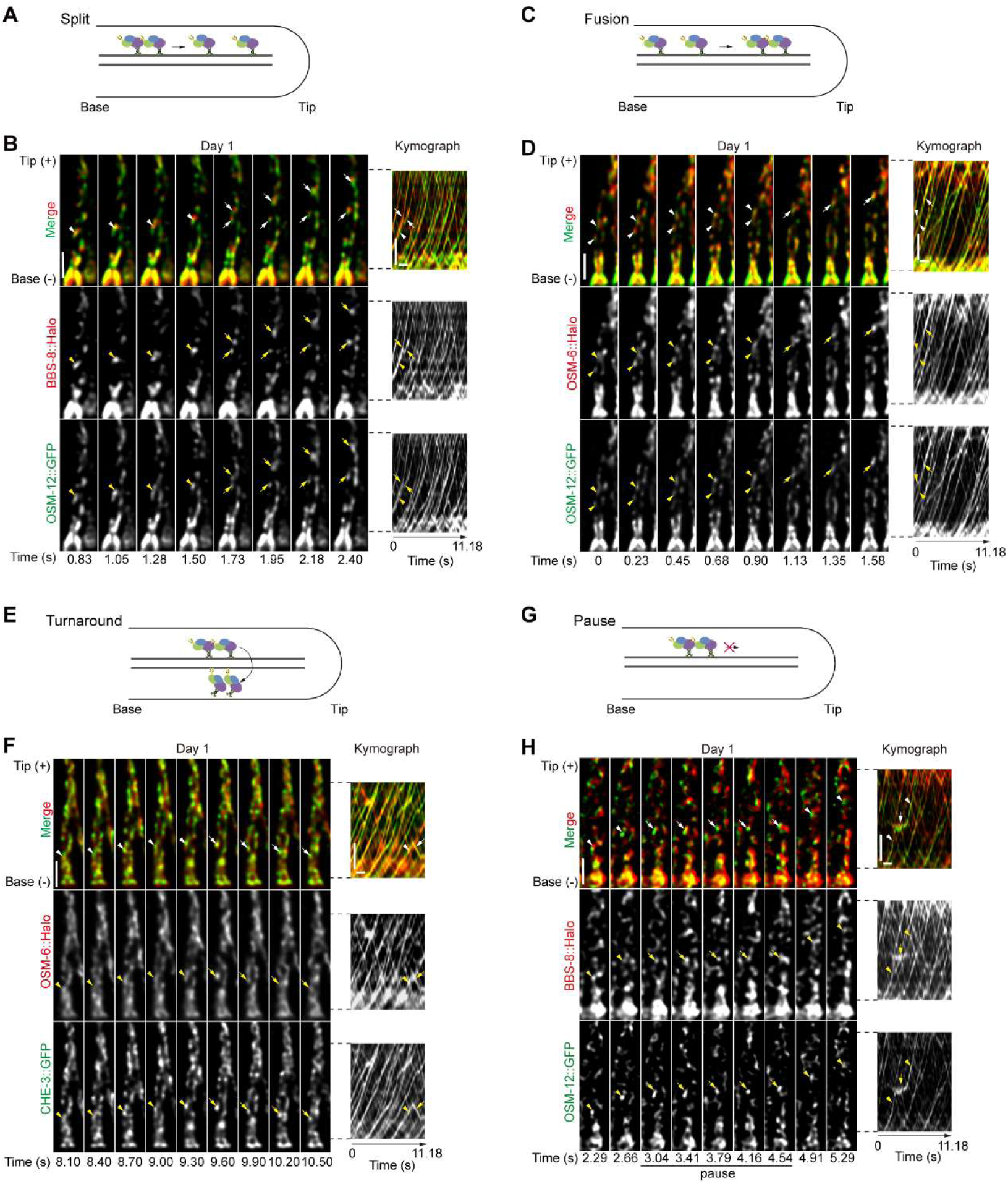
Dynamic behaviors of IFT trains in worm cilia. **(A, C, E**, and **G)** Depictions of the split (A), fusion (C), intraciliary turnaround (E), and pause (G) of IFT trains. **(B, D, F, and H)** Representative time lapse images and corresponding kymographs of the indicated pair of IFT subunits undergoing ‘split’ (B), ‘fusion’ (D), ‘turnaround’ (F), and ‘pause’ (H). Arrowheads denote IFT complexes before the behavioral events, and arrows denote IFT complexes after the behavioral events in (B, D and F). Arrowheads denote IFT complexes before and after the ‘pause’ events, and arrows denote IFT complexes during the ‘pause’ events in (H). Scale bars: 1 μm in time lapse images. In kymographs, the horizontal scale bars: 1 s, the vertical scale bars: 1 μm. Please also see Videos EV2-5.

Taken together, these data further demonstrate the spatial and temporal resolution of our live-imaging analysis and indicate that in addition to the velocity and incidence of IFT trains, ageing is associated with a remarkable change in their behavior as well. These age-dependent changes in the behaviors of IFT trains could underlie the deterioration in ciliary transport with ageing.

### IFT subunits dissociate during ciliary transport

We next examined individual IFT components in our 12 subunit combinations. Although IFT proteins have been considered to move together in a complex (Nakayama & Katoh, 2018), we observed clear dissociations between IFT components (hereafter referred as ‘subunit dissociation’) in all 12 combinations (Fig. 3A,B and Video EV6). Subunit dissociation generated a dual-labeled particle and a mono-labeled particle (Fig. 3A,B and Video EV6), implying that it happened in part of the IFT complexes of an IFT train. Besides, the mono-labeled particle still moved post dissociation (Fig. 3B and Video EV6), indicating that it is associated with the motor and potentially other IFT components as well.

**Figure 3.**
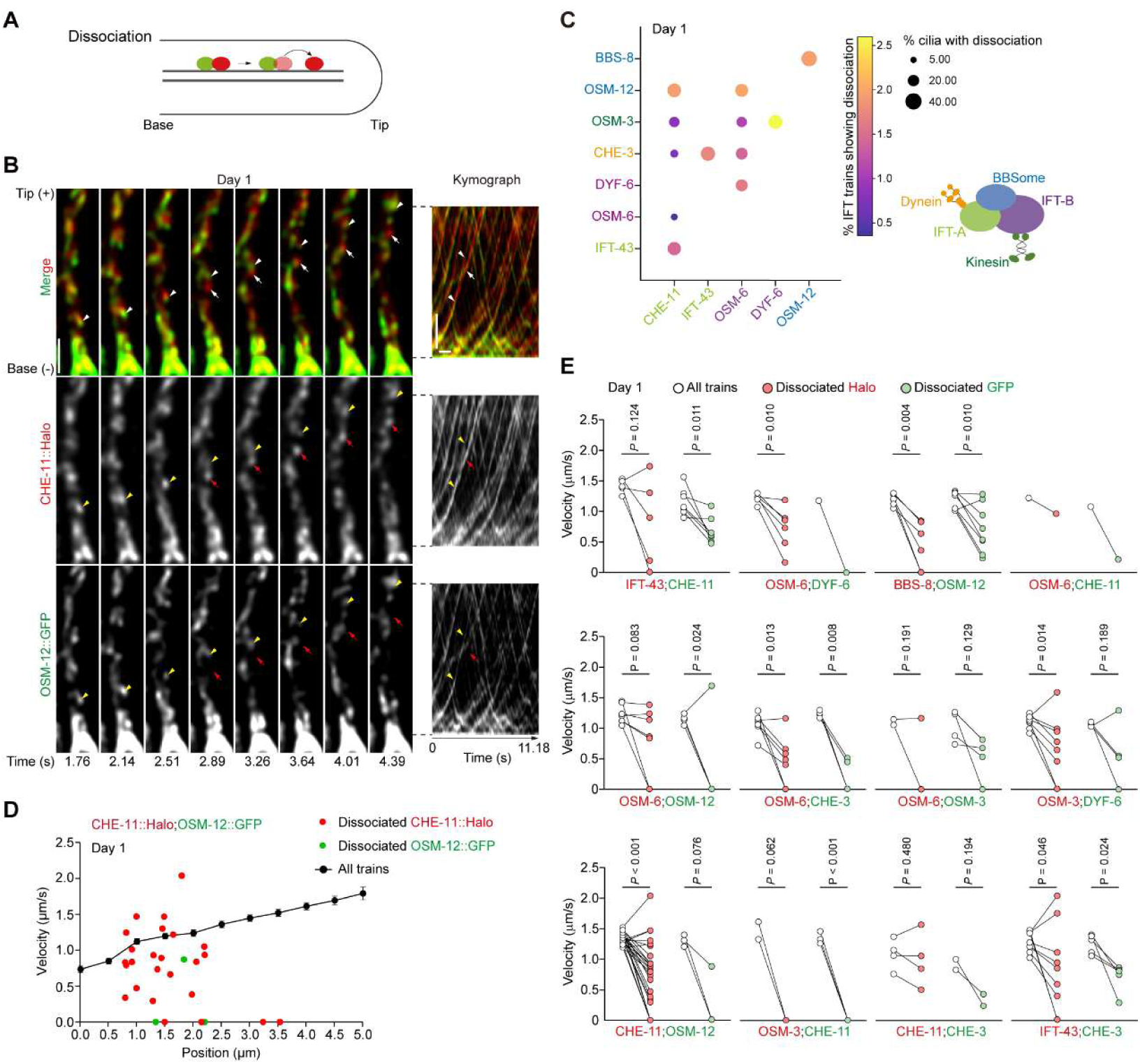
IFT subunits dissociate during ciliary transport. **(A)** A depiction of the dissociation event. **(B)** Representative time lapse images and corresponding kymographs showing the dissociation between CHE-11::Halo and OSM-12::GFP in the sensory cilia of young worms. Arrowheads denote the IFT complex suffering dissociation, arrows denote the dissociated CHE-11::Halo. Scale bars: 1 μm in time lapse images. In kymographs, the horizontal scale bar: 1 s, the vertical scale bar: 1 μm. Please also see Video EV6. **(C)** The incidence of dissociation varies across examined IFT subunit pairs. **(D)** The velocities of dissociated CHE-11::Halo (red dots), OSM-12::GFP (green dots), and all IFT trains (black dots) along the cilia. Error bars: SEM. **(E)** In general, dissociation reduces IFT velocity in the 12 examined subunit pairs. Paired *t*-test.

In healthy young worms, IFT components dissociates with each other at a low incidence (<3% of all examined IFT trains) (Fig. 3C and Fig. EV3A). However, this phenomenon is prevalent in worm cilia. In our 11-sec long observation windows, up to 36.7% cilia showed IFT components dissociation (Fig. 3C and Fig. EV3A). Moreover, the incidence of subunit dissociation varied across the 12 combinations examined (Fig. 3C and Fig. EV3A). For example, the pair of CHE-11, the IFT-A core subunit, and OSM-12, the BBSome component, exhibited a high dissociation incidence among the examined combinations (Fig. 3C and Fig. EV3A), whereas CHE-11 and the IFT-B core component OSM-6 showed the lowest dissociation incidence (Fig. 3C and Fig. EV3A).

We further assessed the functional consequence of subunit dissociation by analyzing IFT velocity. In all examined 12 combinations, mono-labeled IFT trains usually moved slower than the dual-labeled ones (Fig. 3D,E and Fig. EV3B), indicating that the dissociation between IFT components impairs ciliary transport.

### Ageing increases the dissociation between IFT components

We previously reported that ciliary transport decreases with age. Since the dissociation between IFT components reduces IFT velocity, we speculate that subunit dissociation could become more prevalent in aged worms and contribute to the age-dependent deterioration of IFT. In line with our hypothesis, we observed that the incidence of subunit dissociation generally increased in aged worms (Fig. 4A-C, Fig. EV3A and Video EV7). As in young worms, the incidence of subunit dissociation varied in examined pairs (Fig. 4B and Fig. EV3A). Moreover, the age-dependent increase in subunit dissociation also varied in different pairs of IFT components (Fig. 4C). Notably, in the five pairs showing a remarkable age-dependent increase in subunit dissociation (Fig. 4C,D), four were tightly bond in young worms (Fig. 3C). On the contrary, the pairs showing weaker interactions in young worms (e.g., the pair of CHE-11 and OSM-12) did not show a significant increase in subunit dissociation during ageing (Fig. 3C, 4C and 4D). Therefore, ageing is likely to compromise the stronger IFT subunit interactions in young worms.

**Figure 4.**
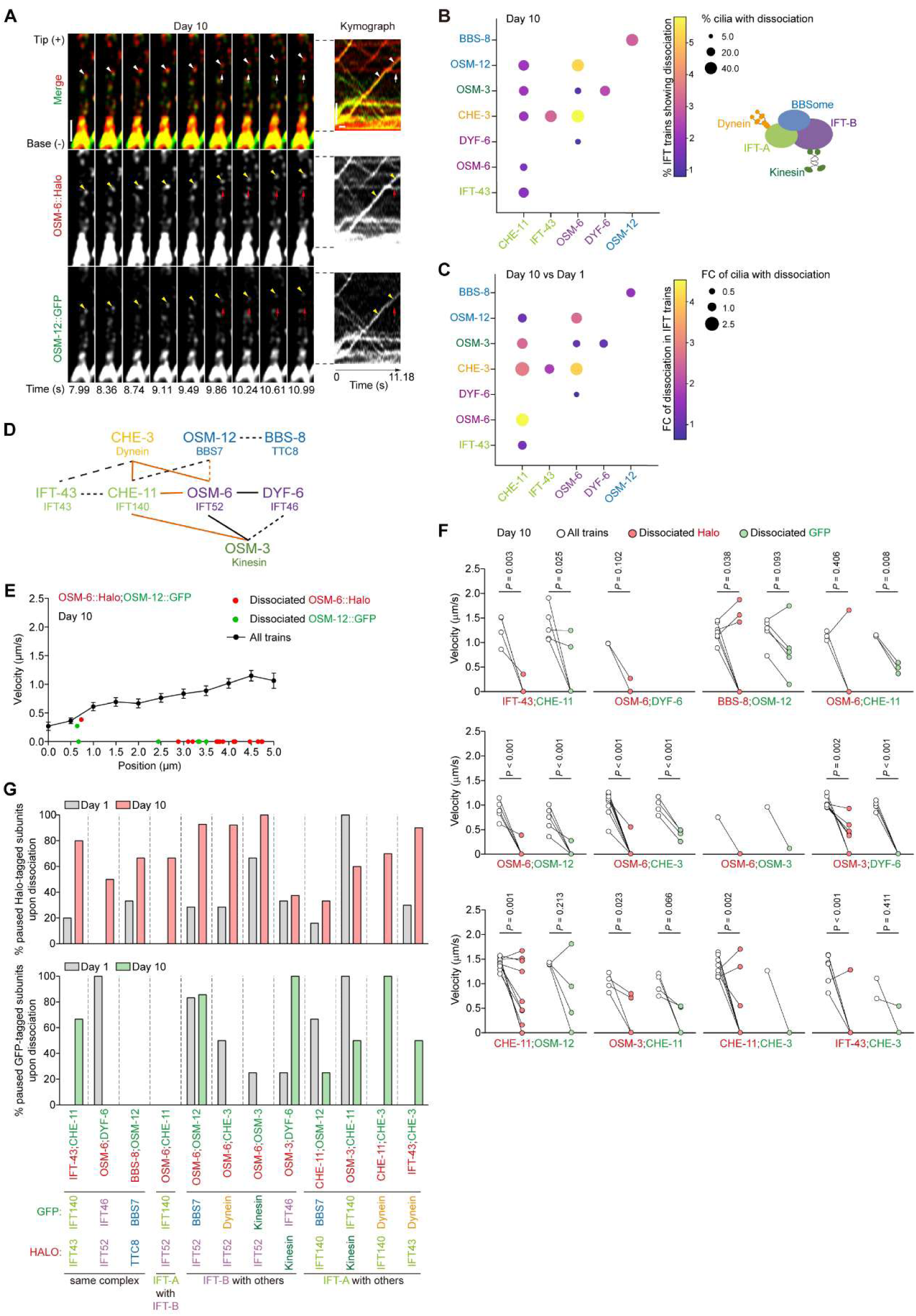
Ageing impairs the integrity of IFT complex selectively. **(A)** Representative time lapse images and corresponding kymograph showing the dissociation between OSM-6::Halo and OSM-12::GFP in the sensory cilia of aged worms (day 10 of adulthood). Arrowheads denote IFT complex suffering dissociation, arrows denote dissociated IFT subunits. Scale bars: 1 μm in time lapse images. In kymographs, the horizontal scale bar: 1 s, the vertical scale bar: 1 μm. Please also see Video EV7. **(B)** The incidence of dissociation in examined IFT subunit pairs at day 10 of adulthood. **(C)** The dissociation of IFT subunits increases during ageing. FC: fold of change. **(D)** A model summarizing the dissociation between indicated IFT subunits. Dashed lines denote the weak interactions in young worms. Orange denotes the age-dependent increase in subunit dissociation. **(E)** The velocities of dissociated OSM-6::Halo and OSM-12::GFP in the sensory cilia of aged worms, in comparison with the mean velocity of all OSM-6::Halo;OSM-12::GFP-labeled trains. Error bars: SEM. **(F)** The comparisons of the velocities of dissociated IFT subunits and corresponding IFT trains in aged worms. Paired *t*-test. Halo-tagged subunits are marked in red, whereas GFP-tagged ones in green. **(G)** The percentages of paused IFT subunits upon dissociation. Note that ageing increases the incidence of pause of dissociated IFT subunits.

We further examined the impact of subunit dissociation on IFT in aged worms. Whereas subunit dissociation also slowed down IFT in aged worms, it caused a bigger reduction in IFT velocity (Fig. 4E,F and Fig. EV3B). Of note, more dissociated IFT subunits were found to ‘pause’ in aged worms (Fig. 4G).

### The molecular chaperone TRiC/CCT regulates the age-dependent instability of IFT complex

We next investigated how ageing increases subunit dissociation. The age-dependent transcriptomic changes in worm neurons showed a significant downregulation of *cct-1* and *cct-8* (Fig. 5A) (Wang *et al*, 2022). The two genes encode subunits of the eukaryotic chaperonin TCP-1 ring complex (TRiC/CCT), a complex responsible for the folding of IFT components (Zhao *et al*, 2026). By qPCR of isolated neurons, we confirmed that the two TRiC/CCT subunits decreased with age in worm neurons (Fig. 5B). Since proper folding is critical to the formation of protein complex, the age-dependent reduction in *cct-1* and *cct-8* could cause subunit dissociation.

**Figure 5.**
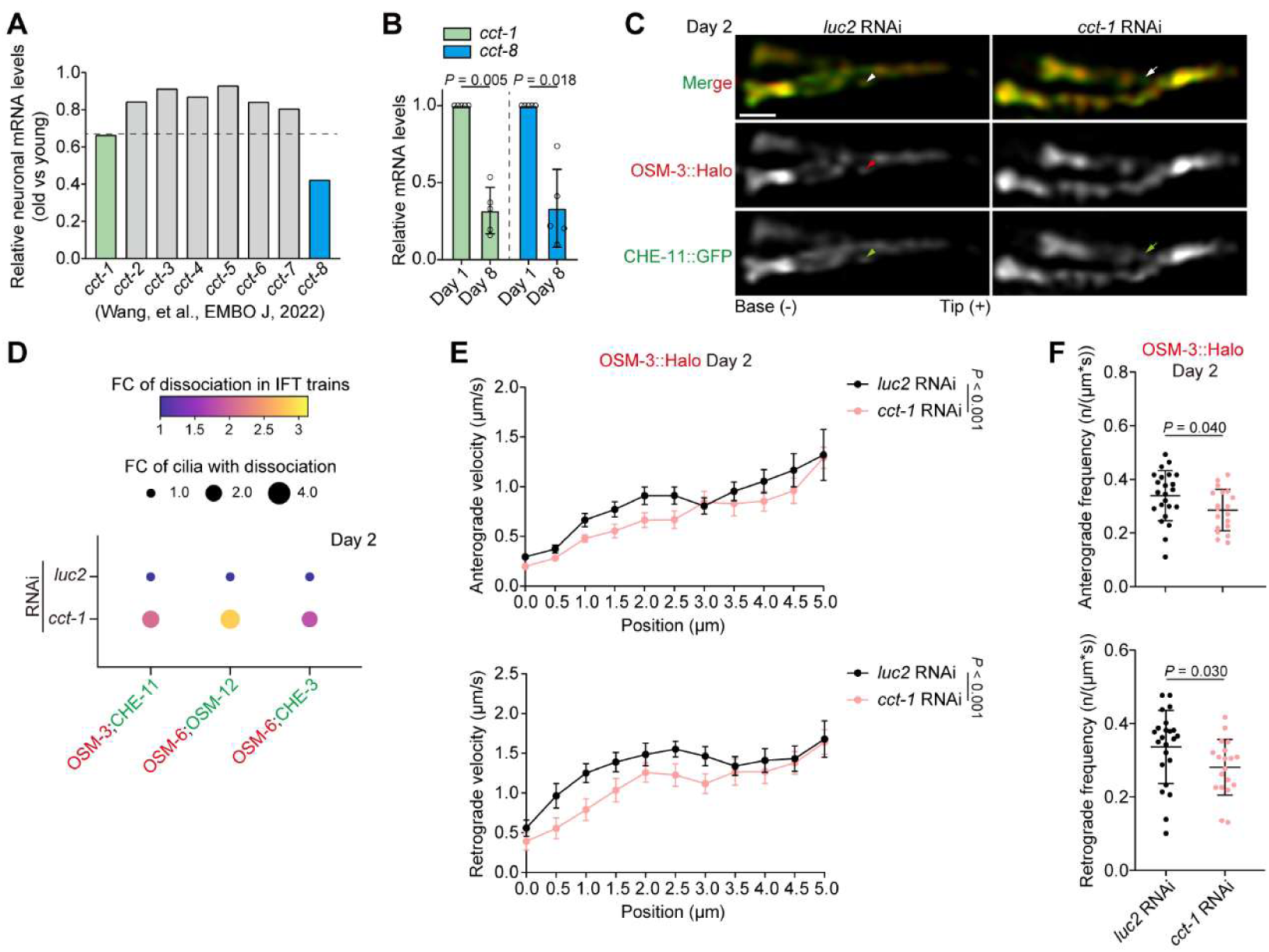
The age-dependent reduction in TRiC complex promoted the dissociation of IFT subunits. **(A)** The transcriptional changes of indicated TRiC complex components. RNA-Seq data from Wang, et al., EMBO J, 2022 (PMID: 35253240). **(B)** qPCR of *cct-1* and *cct-8* in isolated worm neurons at indicated ages. Error bars: SD. Paired *t*-test. **(C)** Representative images of OSM-3::Halo;CHE-11::GFP labeled sensory cilia in young worms upon indicated RNAi treatments. Worms were treated with RNAi from the fourth larval stage to day 2 of adulthood. *luc2*, a firefly luciferase gene irrelevant to *C. elegans* biology, serves as the negative control. Arrows denote dissociated CHE-11::GFP, arrowheads denote IFT subunits with both fluorescence. Scale bar: 1 μm. **(D)** Suppressing *cct-1* in the neurons of young worms increases dissociation of indicated IFT subunit pairs. Halo-tagged subunits are marked in red, whereas GFP-tagged ones in green. FC: fold of change. **(E and F)** Neuron-specific RNAi against *cct-1* in young worms reduced OSM-3::Halo velocities (E) and frequencies (F) in the sensory cilia. OSM-3::Halo is from the sensory cilia expressing both OSM-3::Halo and CHE-11::GFP. Error bars: SEM in (E) and SD in (F). Two-way ANOVA in (E), and Mann-Whitney test in (F).

To examine this hypothesis, we first performed neuron-specific RNAi against *cct-1* in young worms (Fig. EV4A). OSM-3 & CHE-11, OSM-6 & OSM-12, and OSM-6 & CHE-3 are three pairs showing remarkable in subunit dissociation during ageing (Fig. 4C,D). As expected, neuron-specific suppression of *cct-1* increased the dissociation incidence of the three pairs (Fig. 5C,D and Fig. EV4B).

In line with our speculation that the increased subunit dissociation contributes to the impaired IFT in aged worms, neuron-specific RNAi against *cct-1* reduces the velocity and the frequency of OSM-3::Halo in the young worms expressing OSM-3::Halo and CHE-11::GFP (Fig. 5E,F and Video EV8).

To confirm the role of TRiC/CCT in the age-dependent instability of IFT complex, we next upregulated *cct-1* and *cct-8* in the sensory neurons of aged worms (Fig. EV5A). Increasing *cct-1* and *cct-8* reduced the dissociation in the pairs of OSM-3 & CHE-11 and OSM-6 & OSM-12, but not in the pair of OSM-6 & CHE-3 (Fig. EV5B-D), suggesting that TRiC/CCT-mediated protein folding controlled the interactions of different IFT subunits at various levels. Consistent with the reduction of dissociation between OSM-3::Halo and CHE-11::GFP, overexpressing *cct-1* and *cct-8* increased the velocity of OSM-3::Halo (Fig. EV5E and Video EV9). However, the upregulation of the two TRiC/CCT components did not increase the frequency of OSM-3::Halo particles (Fig. EV5F and Video EV9), suggesting other regulators in the age-dependent decrease of IFT.

Taken together, these results indicate that ageing impairs the stability of IFT complex and its movement through the downregulation of the molecular chaperone TRiC/CCT.

### *daf-19* is responsible for the age-dependent instability of IFT complex

The RFX transcription factor, *daf-19*, is the master regulator of IFT genes in *C. elegans* (Senti & Swoboda, 2008). We previously reported that its downregulation during ageing reduces the global level of IFT genes and, in turn, impairs IFT in aged worms (Zhang *et al*., 2021a). The formation of IFT complex requires a stable pool of IFT subunits. Therefore, the age-dependent decrease in IFT genes levels could cause the instability of IFT complex.

To test this hypothesis, we overexpressed *daf-19c*, the *daf-19* isoform which specifically regulates ciliary genes, in sensory neurons and examined the incidence of subunit dissociation in aged worms (Fig. 6A and Fig. EV6A). Indeed, *daf-19c* overexpression significantly reduced dissociation in all three pairs of IFT subunits (i.e., OSM-3 & CHE-11, OSM-6 & OSM-12, and OSM-6 & CHE-3) (Fig. 6B and Fig. EV6B). As we reported (Zhang *et al*., 2021a), the velocity and the frequency of OSM-3::Halo were increased upon *daf-19c* overexpression in the aged worms expressing OSM-3::Halo and CHE-11::GFP (Fig. 6C,D and Video EV10). Therefore, the *daf-19*-controlled expression of IFT components is critical to the age-dependent instability of IFT complex and, in turn, the reduced IFT in aged worms.

**Figure 6.**
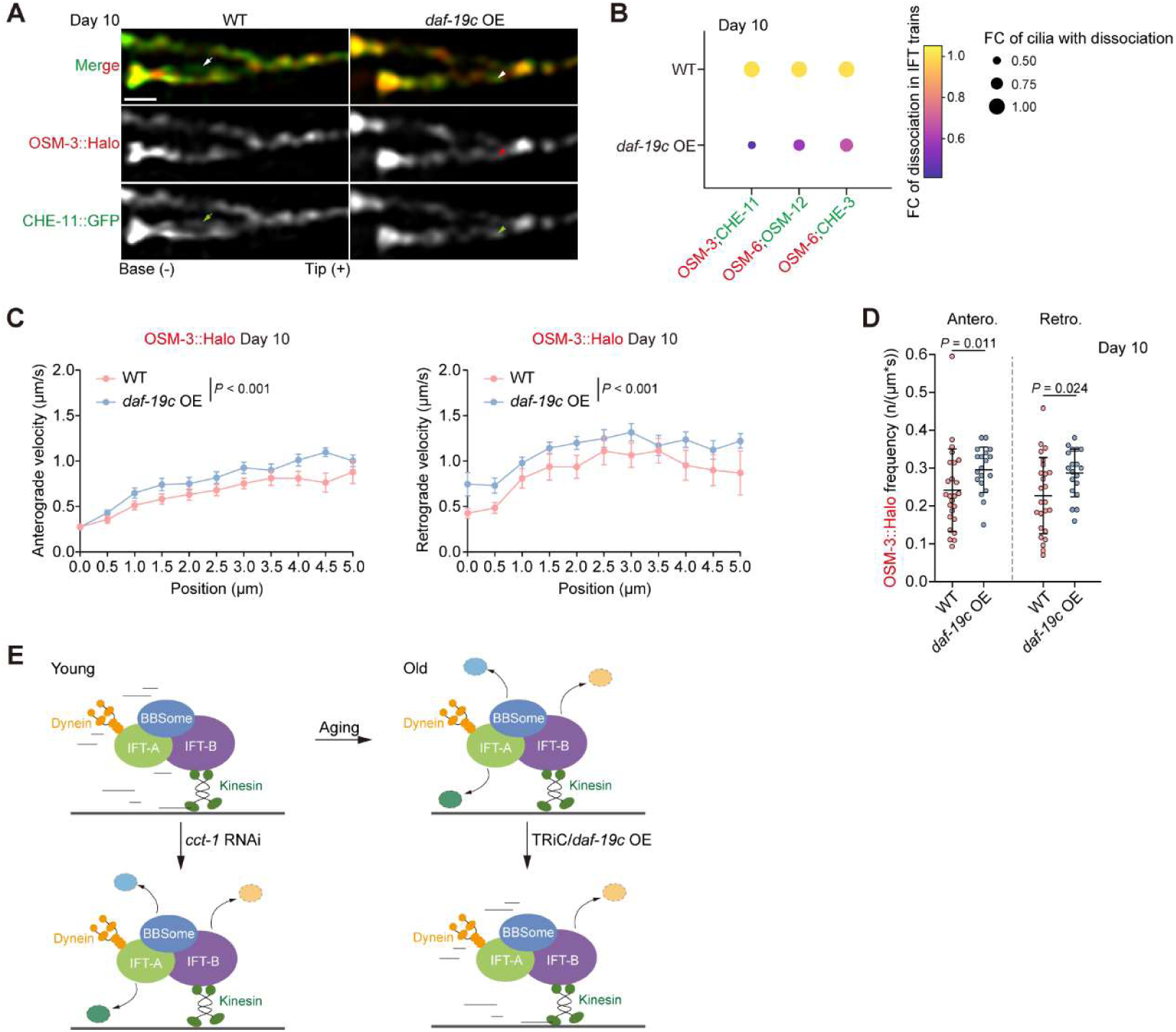
Overexpressing *daf-19c* inhibits the dissociation of IFT subunits in aged worms. **(A)** Representative images of OSM-3::Halo;CHE-11::GFP labeled sensory cilia in aged wild type worms and worms overexpressing *daf-19c*. Arrows denote dissociated CHE-11::GFP, arrowheads denote IFT subunits with both fluorescence. Scale bar: 1 μm. **(B)** Overexpressing *daf-19c* reduces the dissociation of indicated IFT subunits in aged worms at day 10 of adulthood. Halo-tagged subunits are marked in red, whereas GFP-tagged ones in green. FC: fold of change. **(C and D)** Overexpressing *daf-19c* increases the velocity (C) and frequency (D) of OSM-3::Halo in the sensory cilia of aged worms. OSM-3::Halo is from the sensory cilia expressing both OSM-3::Halo and CHE-11::GFP. Two-way ANOVA in (C), Mann-Whitney test in (D). Error bars: SEM in (C) and SD in (D). **(E)** Model. See Discussion for details.

## Discussion

Live tracking of intraflagellar transport (IFT) trains is critical for understanding the molecular mechanics of ciliary transport. Here, we present a comprehensive, super-resolution, live-imaging analysis of IFT in young and aged *C. elegans*. While previous studies have largely tracked single-labeled IFT trains, we multiplexed fluorophores to simultaneously monitor pairs of IFT subunits. This approach revealed that instead of being static entities, IFT complexes undergo highly dynamic compositional changes while traversing the cilium.

The central discovery of our study is the high prevalence of intra-complex dissociation within moving IFT trains. In addition to the known dynamic changes between discrete IFT trains, such as ‘split’ and ‘fusion’ events (Fig. 2), we observed differentially labeled IFT subunits uncoupled during transport in over one-third of the observed cilia (Fig. 3). Given our brief observation window of 11 seconds, it is highly probable that such intra-complex dissociation is a near-ubiquitous feature of the IFT machinery *in vivo*.

Focusing on anterograde transport (Nakayama & Katoh, 2018), we found that this uncoupling extends beyond simple cargo unloading. Dissociation occurred across all 12 examined subunit pairs, including critical interfaces between motor and subcomplexes (e.g., kinesin/IFT-A, dynein/IFT-A) and intra-complex pairings (e.g., BBSome/BBSome). Furthermore, peripheral subunits, such as IFT-43 and specific BBSome components (Nakayama & Katoh, 2018), exhibited higher dissociation frequencies than core components (Fig. 3). The instability observed across all IFT modules suggests that the IFT train undergoes continuous structural remodeling, rather than operating as a rigid vehicle. Given the high prevalence of uncoupling among all 12 labeled pairs, other unlabeled subunits likely experience similar dynamic instability.

Intriguingly, we observed that most dissociated IFT subunits in young worms often retained motility, albeit at significantly reduced velocities (Fig. 3). This suggests that IFT complexes possess a degree of plasticity: either these dissociated fragments retain associated with trace, undetectable amounts of intact IFT trains, or incomplete IFT complexes remain capable of transport provided the overall ciliary architecture is preserved. This latter hypothesis aligns with previous *in vitro* reconstitutions of the *C. elegans* IFT machinery, which demonstrated that partial complexes can indeed translocate, though significantly slower than fully assembled counterpart (Mohamed *et al*, 2018). Our *in vivo* data provide the first physiological corroboration of this modularity, suggesting that the integrity of the IFT machinery directly dictates its transport efficiency.

We observed that ageing markedly exacerbates subunit dissociation, particularly among subunit pairs that remain highly stable in young animals (Fig. 4). This indicates that ageing compromises the core structural integrity of the IFT machinery. Consequently, dissociation in aged worms corresponds with a more severe reduction in IFT velocity (Fig. 4), potentially due to the cumulative loss of multiple, unlabeled stabilizing subunits. This structural deterioration likely represents a primary driver of the age-dependent decline in IFT efficacy and possibly ciliary signaling.

Mechanistically, our data link the age-dependent structural instability to the decline of two key regulatory factors, precise protein folding and stoichiometric supply of its constituent subunits (Fig. 6E). We identified the conserved TRiC/CCT chaperonin complex, which folds IFT proteins (Zhao *et al*., 2026), as an essential regulator of IFT subunit stability. The age-dependent downregulation of TRiC/CCT leads to increased uncoupling, while its upregulation partially rescues complex integrity and transport velocity in aged worms. Furthermore, we show that upregulating *daf-19*/RFX, the master transcription factor of IFT genes (Senti & Swoboda, 2008; Zhang *et al*., 2021a), mitigates the age-dependent IFT dissociation, underscoring the importance of subunit stoichiometry for complex stability. While enhancing both folding and supply only partially ameliorates the aged IFT phenotype, the age-related deterioration of IFT is likely governed by multiple converging mechanisms, perhaps including a broader collapse of the proteostasis network. Identifying these additional determinants remains a critical avenue for future research.

In conclusion, our study re-frames the IFT train as a dynamic assembly whose structural integrity is actively maintained. The identification of progressive intra-complex dissociation as a driver of ciliary dysfunction provides a new understanding of both natural ageing and the pathogenesis of late-onset ciliopathies.

## Materials and Methods

### Reagents and tools table

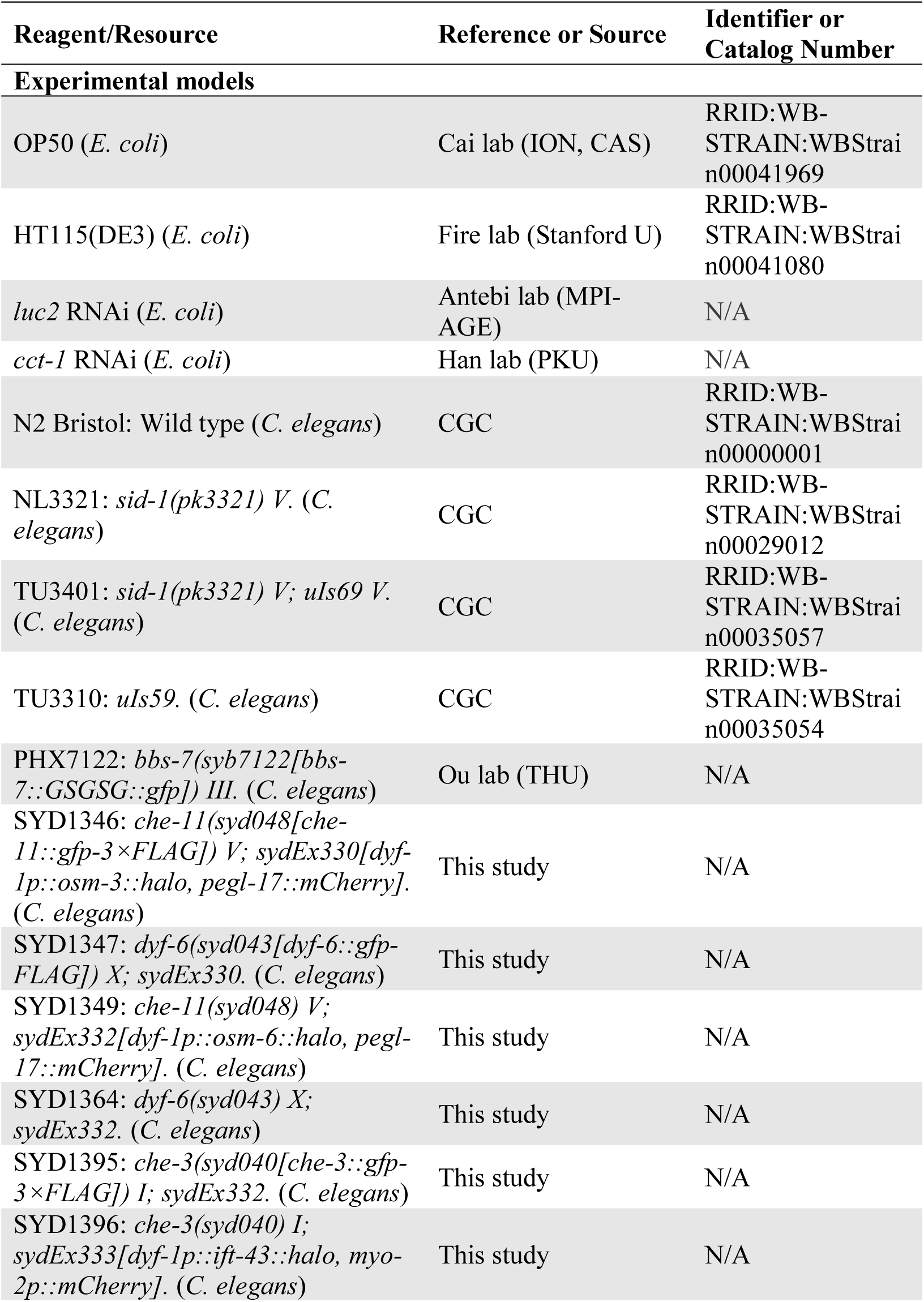
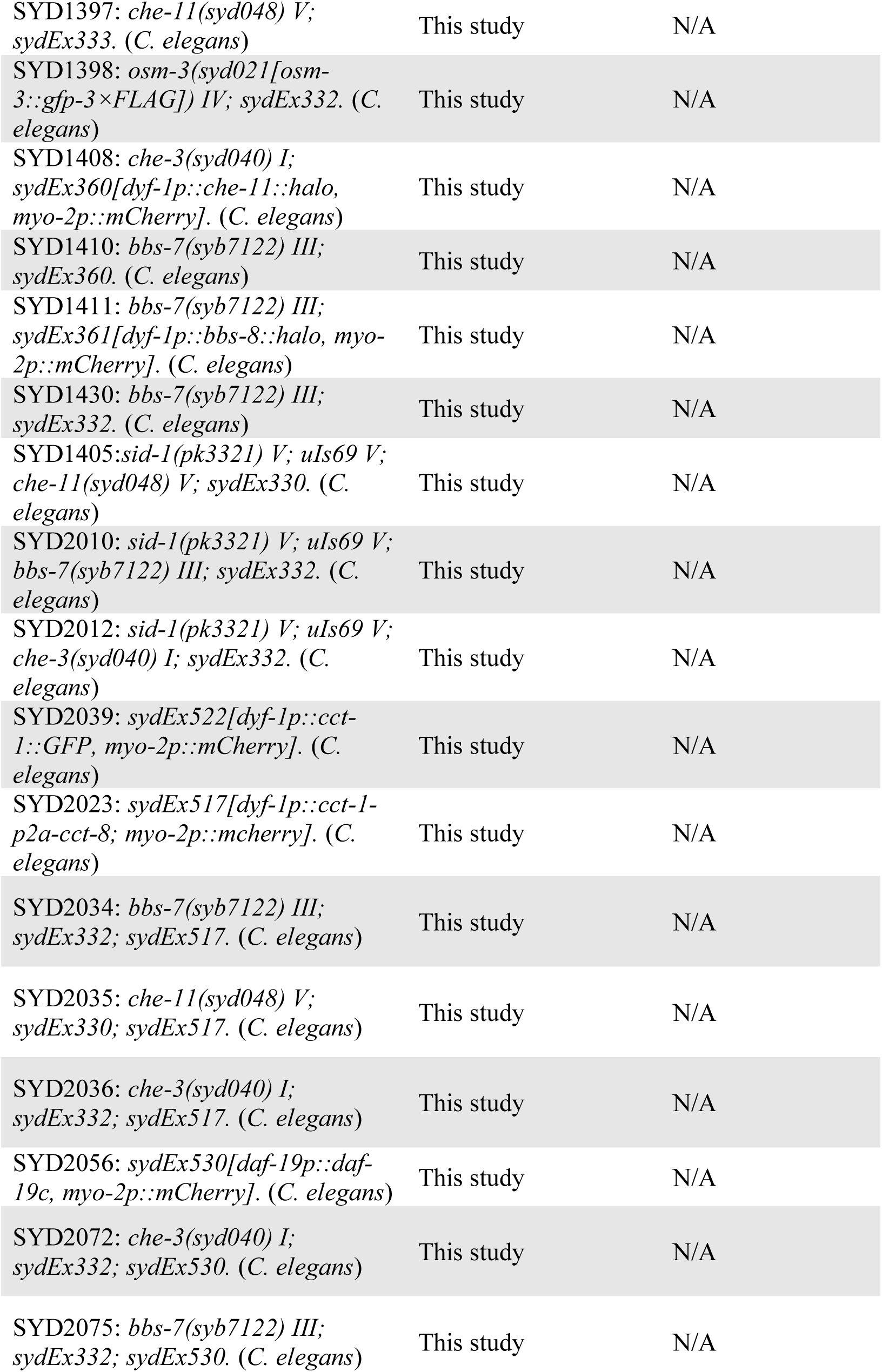
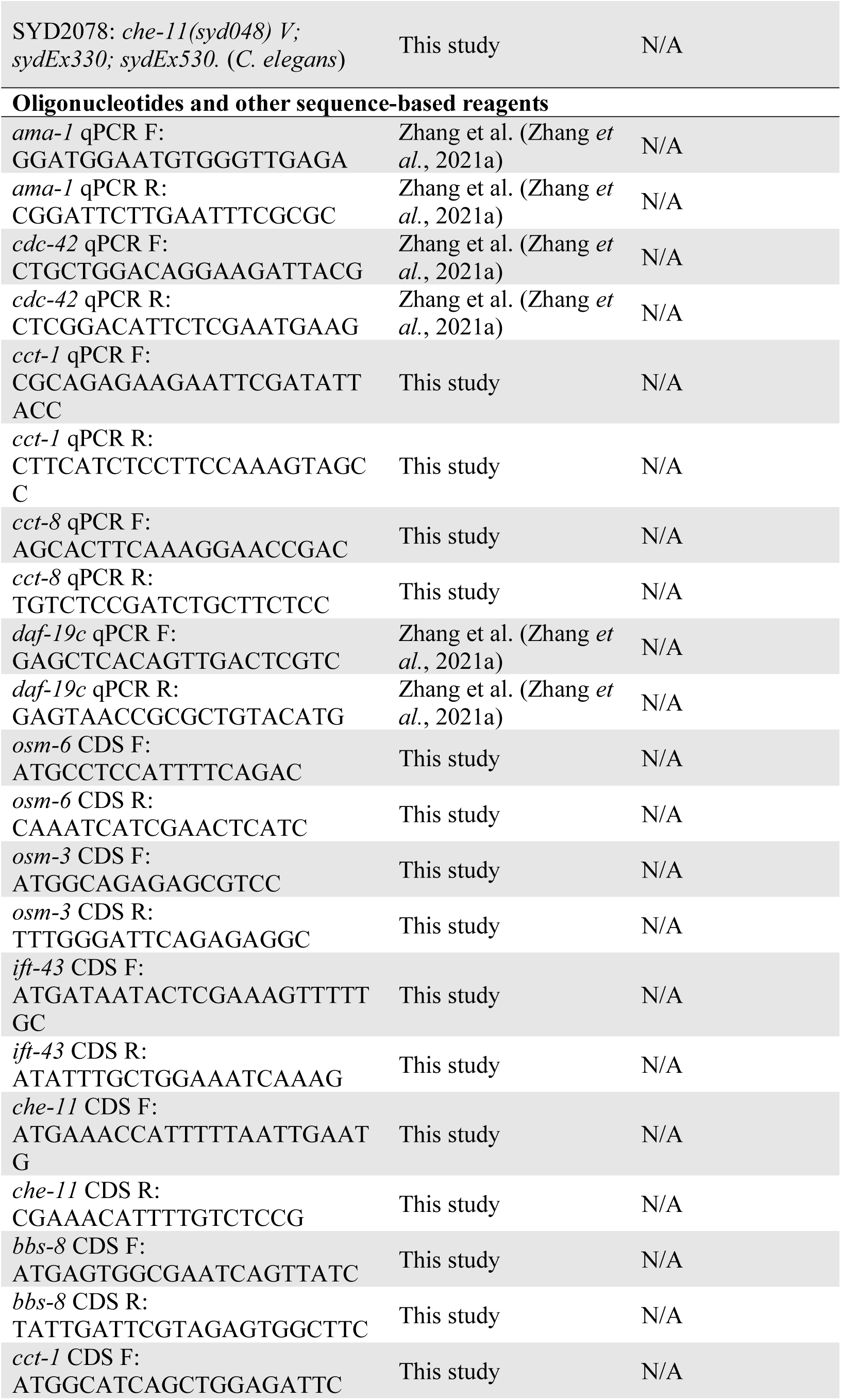
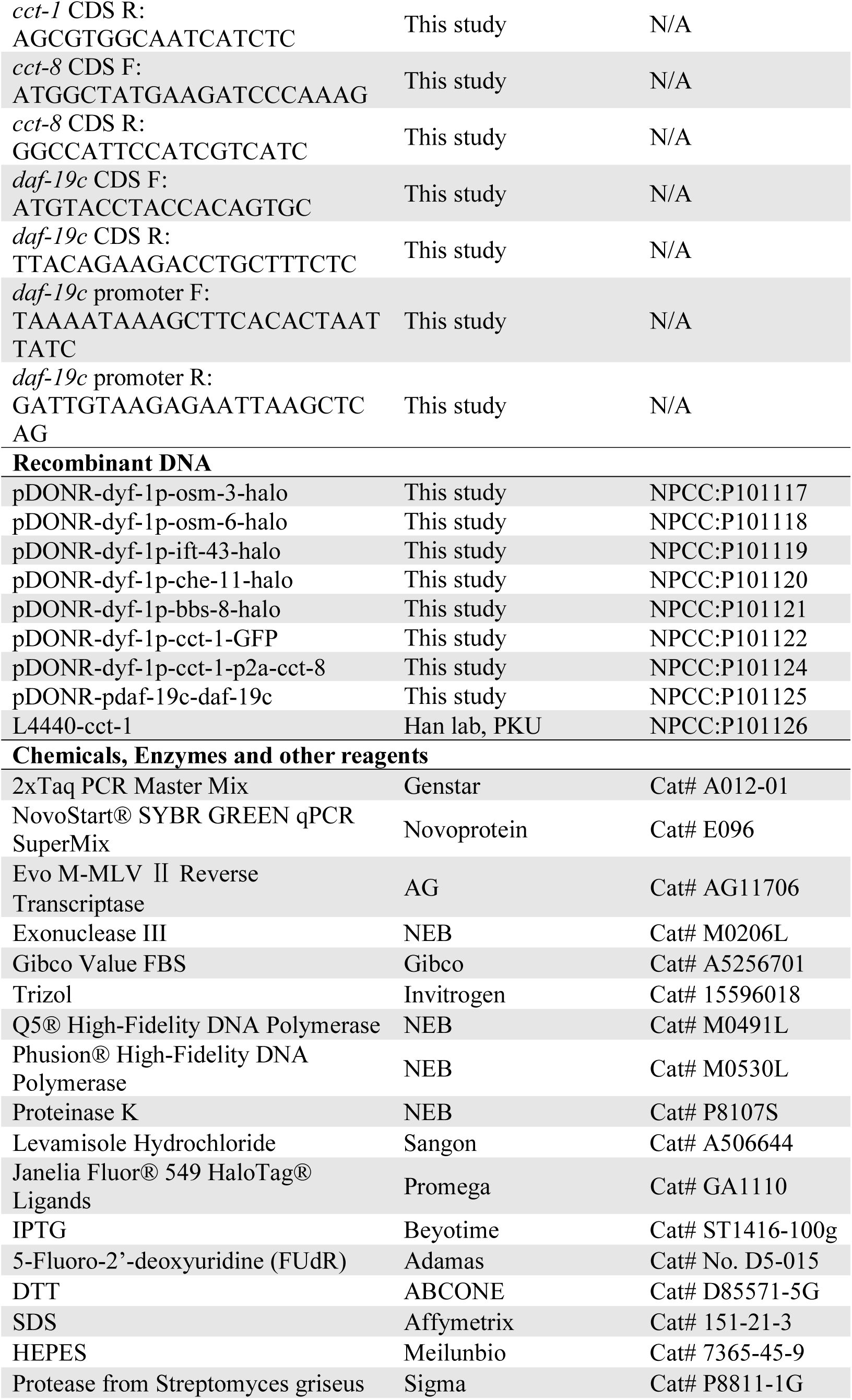
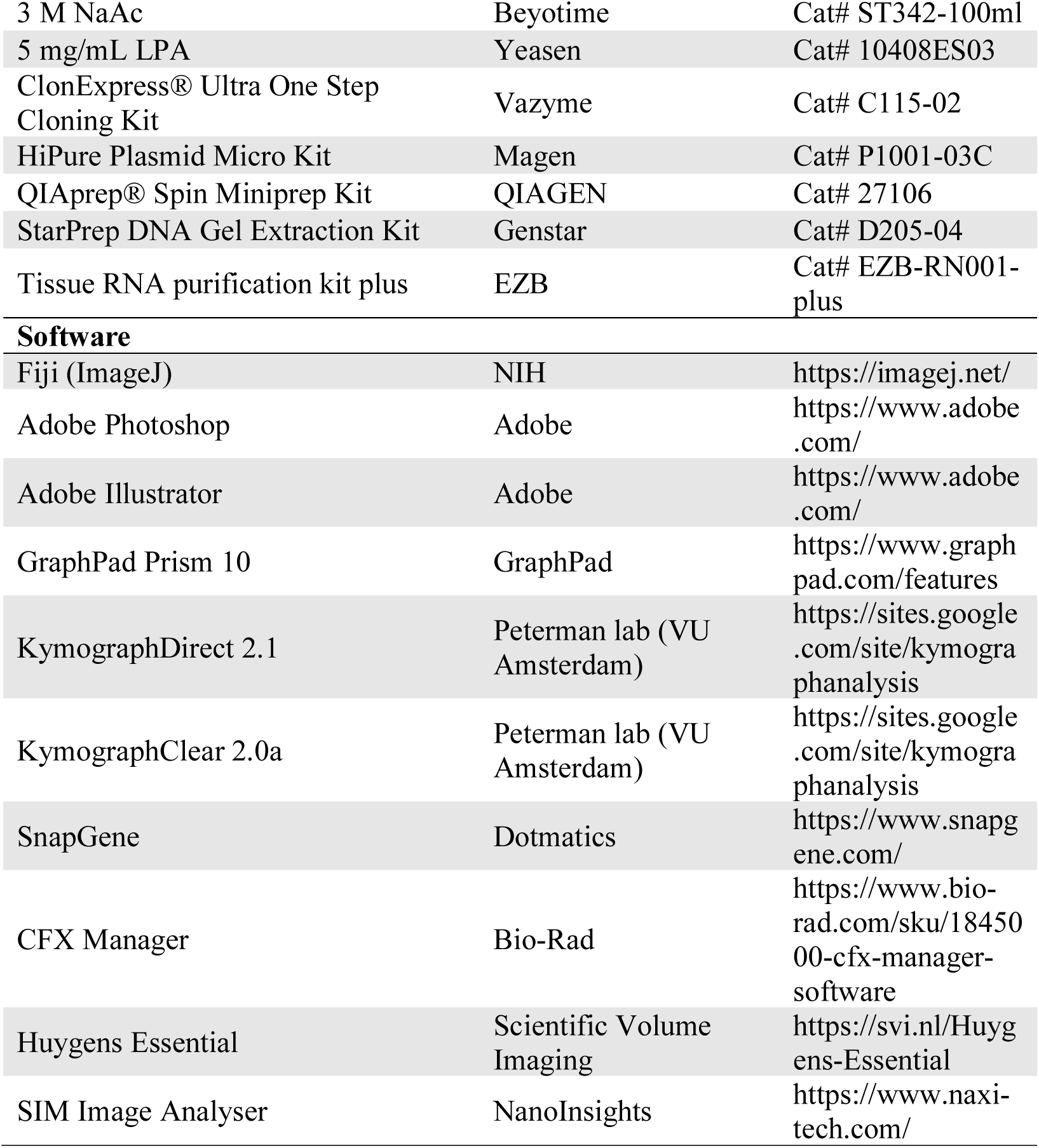

### *C. elegans* strains and culture

*C. elegans* strains used in this study are listed in reagents and tools table. Worms were grown on NGM plates with standard techniques at 20°C (Brenner, 1974). All assayed worms were at day 1 of adulthood unless otherwise noted. Some strains were provided by CGC, which is funded by NIH Office of Research Infrastructure Programs (P40 OD010440). OP50 colonies on an NGM plate were treated with the sterilisation programme in a GS Gene Linker UV Chamber (BIO-RAD) for 15 min to obtain UV-killed bacteria. The killed bacteria were confirmed by no growth after O/N incubation in LB.

To isolate worm neurons, ∼8,000 synchronised TU3310 worms of day 1 adulthood were collected. For day 8 samples, ∼16,000 synchronised TU3310 L4 larvae were transferred onto plates supplemented with 25 μM FUdR (Adamas) (Gandhi *et al*, 1980; Mitchell *et al*, 1979). Worms were washed three times with M9 to remove remaining bacteria and then transferred onto fresh NGM plates with FUdR every other day from L4 to day 8 of adulthood.

### Plasmid

All plasmids used in this study were validated by sequencing and deposited into National Plasmid Collection Center (NPCC), China. A complete list of plasmids and their deposit numbers is provided in reagents and tools table.

### Transgenes

Extra-chromosomal transgenic lines were obtained by co-injecting the plasmids of *dyf-1p::osm-3::halo* and *egl-17p::mCherry*, *dyf-1p::osm-6::halo* and *egl-17p::mCherry*, *dyf-1p::ift-43::halo* and *myo-2p::mCherry*, *dyf-1p::che-11::halo* and *myo-2p::mCherry*, *dyf-1p::bbs-8::halo* and *myo-2p::mCherry*, *dyf-1p::cct-1::gfp* and *myo-2p::mCherry*, *dyf-1p::cct-1::p2a::cct-8* and *myo-2p::mCherry*, *daf-19p::daf-19c* and *myo-2p::mCherry* into N2. Plasmid concentrations for microinjections were 50 ng/μl for the genes of interest, 20 ng/μl for the injection marker *egl-17p::mCherry*, and 2.5 ng/μl for the injection marker *myo-2p::mCherry*.

For knock-in lines, injections and subsequent screens were performed as described (Dickinson *et al*, 2015). Self-excising selection cassettes were discarded before sequencing and phenotypic analysis. All knock-in strains were verified by DNA sequencing.

### HaloTag staining

30-40 L4 worms were stained in 50 μl S-medium with concentrated OP50 bacteria and 2.5 μM of JF549 HaloTag ligand (Promega) in a darkened 96-well plate shaking at 150 rpm for 19 hr at 23°C (Grimm *et al*, 2015). Animals were recovered on NGM plates for up to two hours before imaging (Fan *et al*, 2020).

### Microscopy

Worms were anaesthetised with 0.3% levamisole (Sangon) in M9 buffer, mounted on 5% agar pads, and maintained at room temperature. Cilia with the base, proximal and distal segments in focus were examined. To track dual-fluorescence labeled IFT trains in wild type worms, time lapse images of 100 frames were collected using the GI-SIM microscope equipped with a 100×, 1.5 NA objective (Li lab) (Guo *et al*, 2018; Qiao *et al*., 2023). For each frame, 9 images of 3 orientations and 3 phases were captured for each channel with an exposure speed at 2 ms per image. To monitor dual-fluorescence labeled IFT trains in RNAi-treated worms, the worms overexpressing *daf-19c*, or the worms overexpressing *cct-1* and *cct-8*, time lapse images of 100 frames were collected using the Multi-SIM microscope equipped with a 100×, 1.49 NA objective (NanoInsights). For each frame, 15 images of 3 orientations and 5 phases were captured for each channel with an exposure speed at 1 ms per image. Interleaved reconstruction was performed using the reported SIMILR method (Ma *et al*, 2018). Image deconvolution was performed with Huygens Essential (version 23.10, Scientific Volume Imaging, Hilversum, The Netherlands). For other assays, worms were imaged on an Olympus BX53 microscope equipped with a 60×, 1.35 NA objective (Olympus). Fluorescent intensities were measured by ImageJ (NIH).

### Kymograph generation and analysis

Kymographs were generated and analysed as described (Mangeol *et al*, 2016; Prevo *et al*, 2015). In brief, Fourier filtered and separated anterograde or retrograde kymographs were generated with the KymographClear toolset plugin in ImageJ. IFT velocities at every 0.5 μm along cilia were measured by KymographDirect from the kymographs.

### Isolation of worm neurons

Worm neurons were isolated as reported (Kaletsky *et al*, 2015; Wang *et al*., 2022). Worms with neuron-specific fluorescent marker *unc-119p::YFP* (TU3310) were dissociated by SDS–DTT treatment and proteolysis with mechanical disruption. Dissociated worms were filtered with 5 μm cell strainer and subjected to FACS to sort out fluorescent neurons. N2 worms at corresponding age were used as negative control to eliminate auto-fluorescence. Around 8,000 neurons were collected for each biological replicate.

### RNA preparation

To collect total RNA from whole worms, more than 100 synchronized worms were collected into TRIzol (Invitrogen) in each sample. Total RNA was column purified by Tissue RNA purification kit plus (EZB) following the manufacturer’s instruction(Shen *et al*, 2012).

RNA from FACS-sorted neurons was isolated as reported (Wang *et al*., 2022). In brief, the isolated neurons were collected in TRIzol and mixed with chloroform. After centrifugation, the upper aqueous phase was collected, and mixed with a same volume of isopropanol, 0.1 volume of 3 M NaAc, and 3 μl of 5 mg/ml LPA. The mixture was then placed at −20°C to precipitate overnight. The precipitated RNA was washed twice with 75% ethanol, then dissolved in RNase-free water.

### RT-qPCR

cDNA was generated by Evo M-MLV Ⅱ Reverse Transcriptase (AG) for RT-qPCR (Bio-Rad) following the manufacturer’s instruction. qPCR was performed with 2x NovoStart® SYBR GREEN qPCR SuperMix (Novoprotein) on a CFX384 Touch^TM^ Real-Time PCR Detection System (Bio-Rad). A combination of *ama-1* and *cdc-42* was used as reference. At least three technical replicates were performed in each reaction. Primer sequences are listed in reagents and tools table.

### RNA interference

RNAi were performed by feeding worms with *E. coli* HT115 bacteria expressing corresponding dsRNA on standard NGM plates containing 100 μg/ml ampicillin and 1 mM IPTG from the fourth larval stage to day 1 of adulthood, as described (Kamath *et al*, 2001). The worms at day 1 of adulthood were then cultured in liquid RNAi medium for 19 h or overnight. The liquid medium, S-medium, contained 125 mg/ml *E. coli* HT115 bacteria and 1 mM IPTG. HT115 expressing dsRNA against *luc2*, a firefly gene irrelevant to *C. elegans* biology, served as negative control to minimise off-target phenotypes. HT115 strains [*L4440::luc2*] and [*L4440::cct-1*] were kind gifts from Antebi lab (MPI-AGE) and Han lab (Peking U), respectively. For neuron-specific RNAi, the strain TU3401 was used.

### Statistical analysis

Results are presented as Mean ± SD unless otherwise noted. Statistical tests were performed as indicated using GraphPad Prism (GraphPad Software). Detailed statistical information is shown in Table EV1.

## Data availability

The representative images and videos, and detailed statistics are included in the manuscript.

The videos of all behaviors of the 12 IFT combinations are deposited at Mendeley Data with the private link for reviewers: https://data.mendeley.com/preview/2x9zth6jbk?a=633267f4-587d-4ddc-a7e4-696f5acb3c8e.

The dataset will be made publicly available as of the date of publication.

All other data are available from the corresponding authors upon reasonable request.

## Acknowledgements

The authors thank Drs Guangshuo Ou (Tsinghua U) and Jing-Dong J. Han (Peking U) for reagents, Ms Chenxiao Guo (NanoInsights) and the institutional core facilities for cell biology and molecular biology for instrumental and technical support. This research was supported by the Strategic Priority Research Program of the Chinese Academy of Sciences (XDB0990000) and National Natural Science Foundation of China (32470724). Yan X. is supported by Shanghai Leading Talent Program of Eastern Talent Plan.

## Author Contributions

Q.L., D.L., X.Y., and Y.S. conceived the project, designed the experiments, analyzed data, and wrote the manuscript. J.L., M.C., M.S., and X.Z. performed the experiments and analyzed data. S.W. and S.X. performed GI-SIM imaging under the supervision of D.L.. J.T. performed image deconvolution. All authors contributed to manuscript editing.

## References

Bertiaux E, Mallet A, Fort C, Blisnick T, Bonnefoy S, Jung J, Lemos M, Marco S, Vaughan S, Trépout S et al (2018) Bidirectional intraflagellar transport is restricted to two sets of microtubule doublets in the trypanosome flagellum. J Cell Biol 217: 4284–4297

Brenner S (1974) The genetics of Caenorhabditis elegans. Genetics 77: 71–94

Buisson J, Chenouard N, Lagache T, Blisnick T, Olivo-Marin JC, Bastin P (2013) Intraflagellar transport proteins cycle between the flagellum and its base. Journal of cell science 126: 327–338

Chien A, Shih SM, Bower R, Tritschler D, Porter ME, Yildiz A (2017) Dynamics of the IFT machinery at the ciliary tip. eLife 6

Cole DG, Snell WJ (2009) SnapShot: Intraflagellar transport. Cell 137: 784–784 e781

Dentler W (2005) Intraflagellar transport (IFT) during assembly and disassembly of Chlamydomonas flagella. The Journal of cell biology 170: 649–659

Dickinson DJ, Pani AM, Heppert JK, Higgins CD, Goldstein B (2015) Streamlined Genome Engineering with a Self-Excising Drug Selection Cassette. Genetics 200: 1035–1049

Fan X, De Henau S, Feinstein J, Miller SI, Han B, Frøkjær-Jensen C, Griffin EE (2020) SapTrap Assembly of Caenorhabditis elegans MosSCI Transgene Vectors. G3 (Bethesda) 10: 635–644

Gandhi S, Santelli J, Mitchell DH, Stiles JW, Sanadi DR (1980) A simple method for maintaining large, aging populations of Caenorhabditis elegans. Mech Ageing Dev 12: 137–150

Goetz SC, Anderson KV (2010) The primary cilium: a signalling centre during vertebrate development. Nat Rev Genet 11: 331–344

Gopalakrishnan J, Feistel K, Friedrich BM, Grapin-Botton A, Jurisch-Yaksi N, Mass E, Mick DU, Müller RU, May-Simera H, Schermer B et al (2023) Emerging principles of primary cilia dynamics in controlling tissue organization and function. Embo j 42: e113891

Green JA, Mykytyn K (2010) Neuronal ciliary signaling in homeostasis and disease. Cell Mol Life Sci 67: 3287–3297

Grimm JB, English BP, Chen J, Slaughter JP, Zhang Z, Revyakin A, Patel R, Macklin JJ, Normanno D, Singer RH et al (2015) A general method to improve fluorophores for live-cell and single-molecule microscopy. Nat Methods 12: 244–250, 243 p following 250

Guo Y, Li D, Zhang S, Yang Y, Liu JJ, Wang X, Liu C, Milkie DE, Moore RP, Tulu US et al (2018) Visualizing Intracellular Organelle and Cytoskeletal Interactions at Nanoscale Resolution on Millisecond Timescales. Cell 175: 1430–1442.e1417

Hao L, Scholey JM (2009) Intraflagellar transport at a glance. J Cell Sci 122: 889–892

Huang X, Fan J, Li L, Liu H, Wu R, Wu Y, Wei L, Mao H, Lal A, Xi P et al (2018) Fast, long-term, super-resolution imaging with Hessian structured illumination microscopy. Nature biotechnology 36: 451–459

Kaletsky R, Lakhina V, Arey R, Williams A, Landis J, Ashraf J, Murphy CT (2015) The C. elegans adult neuronal IIS/FOXO transcriptome reveals adult phenotype regulators. Nature

Kamath RS, Martinez-Campos M, Zipperlen P, Fraser AG, Ahringer J (2001) Effectiveness of specific RNA-mediated interference through ingested double-stranded RNA in Caenorhabditis elegans. Genome biology 2: RESEARCH0002

Kozminski KG, Johnson KA, Forscher P, Rosenbaum JL (1993) A motility in the eukaryotic flagellum unrelated to flagellar beating. Proc Natl Acad Sci U S A 90: 5519–5523

Lacey SE, Foster HE, Pigino G (2023) The molecular structure of IFT-A and IFT-B in anterograde intraflagellar transport trains. Nat Struct Mol Biol 30: 584–593

Lacey SE, Pigino G (2024) The intraflagellar transport cycle. Nat Rev Mol Cell Biol

Ma Y, Li D, Smith ZJ, Li D, Chu K (2018) Structured illumination microscopy with interleaved reconstruction (SIMILR). J Biophotonics 11

Mangeol P, Prevo B, Peterman EJ (2016) KymographClear and KymographDirect: two tools for the automated quantitative analysis of molecular and cellular dynamics using kymographs. Molecular biology of the cell 27: 1948–1957

Meleppattu S, Zhou H, Dai J, Gui M, Brown A (2022) Mechanism of IFT-A polymerization into trains for ciliary transport. Cell 185: 4986–4998.e4912

Mijalkovic J, Prevo B, Oswald F, Mangeol P, Peterman EJ (2017) Ensemble and single-molecule dynamics of IFT dynein in Caenorhabditis elegans cilia. Nature communications 8: 14591

Mitchell DH, Stiles JW, Santelli J, Sanadi DR (1979) Synchronous growth and aging of Caenorhabditis elegans in the presence of fluorodeoxyuridine. J Gerontol 34: 28–36

Mitra A, Loseva E, Peterman EJG (2024) IFT cargo and motors associate sequentially with IFT trains to enter cilia of C. elegans. Nat Commun 15: 3456

Mohamed MAA, Stepp WL, Okten Z (2018) Reconstitution reveals motor activation for intraflagellar transport. Nature

Mul W, Mitra A, Peterman EJG (2022) Mechanisms of Regulation in Intraflagellar Transport. Cells 11

Nakayama K, Katoh Y (2018) Ciliary protein trafficking mediated by IFT and BBSome complexes with the aid of kinesin-2 and dynein-2 motors. J Biochem 163: 155–164

Ou G, E. Blacque O, Snow JJ, Leroux MR, Scholey JM (2005) Functional coordination of intraflagellar transport motors. Nature 436: 583–587

Oya M, Miyasaka Y, Nakamura Y, Tanaka M, Suganami T, Mashimo T, Nakamura K (2024) Age-related ciliopathy: Obesogenic shortening of melanocortin-4 receptor-bearing neuronal primary cilia. Cell Metab 36: 1044–1058.e1010

Pigino G (2021) Intraflagellar transport. Curr Biol 31: R530–r536

Prevo B, Mangeol P, Oswald F, Scholey JM, Peterman EJ (2015) Functional differentiation of cooperating kinesin-2 motors orchestrates cargo import and transport in C. elegans cilia. Nat Cell Biol 17: 1536–1545

Prevo B, Scholey JM, Peterman EJG (2017) Intraflagellar transport: mechanisms of motor action, cooperation, and cargo delivery. Febs j 284: 2905–2931

Qiao C, Li D, Liu Y, Zhang S, Liu K, Liu C, Guo Y, Jiang T, Fang C, Li N et al (2023) Rationalized deep learning super-resolution microscopy for sustained live imaging of rapid subcellular processes. Nature biotechnology 41: 367–377

Rosenbaum JL, Witman GB (2002) Intraflagellar transport. Nat Rev Mol Cell Biol 3: 813–825

Sanders AA, Kennedy J, Blacque OE (2015) Image analysis of Caenorhabditis elegans ciliary transition zone structure, ultrastructure, molecular composition, and function. Methods Cell Biol 127: 323–347

Senti G, Swoboda P (2008) Distinct isoforms of the RFX transcription factor DAF-19 regulate ciliogenesis and maintenance of synaptic activity. Molecular biology of the cell 19: 5517–5528

Shen Y, Wollam J, Magner D, Karalay O, Antebi A (2012) A steroid receptor-microRNA switch regulates life span in response to signals from the gonad. Science 338: 1472–1476

Silva DF, Cavadas C (2023) Primary cilia shape hallmarks of health and aging. Trends Mol Med 29: 567–579

Singh SK, Gui M, Koh F, Yip MC, Brown A (2020) Structure and activation mechanism of the BBSome membrane protein trafficking complex. Elife 9

Stepanek L, Pigino G (2016) Microtubule doublets are double-track railways for intraflagellar transport trains. Science 352: 721–724

Sun S, Liang B, Koplas A, Tikhonenko I, Nachury M, Khodjakov A, Sui H (2025) Intraflagellar transport trains can switch rails and move along multiple microtubules in intact primary cilia. Proceedings of the National Academy of Sciences of the United States of America 122: e2413968122

Tobin JL, Beales PL (2009) The nonmotile ciliopathies. Genet Med 11: 386–402

Volos P, Fujise K, Rafiq NM (2025) Roles for primary cilia in synapses and neurological disorders. Trends Cell Biol 35: 6–10

Wang X, Jiang Q, Song Y, He Z, Zhang H, Song M, Zhang X, Dai Y, Karalay O, Dieterich C et al (2022) Ageing induces tissue-specific transcriptomic changes in Caenorhabditis elegans. The EMBO journal 41: e109633

Xie C, Li L, Li M, Shao W, Zuo Q, Huang X, Chen R, Li W, Brunnbauer M, Okten Z et al (2020) Optimal sidestepping of intraflagellar transport kinesins regulates structure and function of sensory cilia. The EMBO journal 39: e103955

Zhang Y, Zhang X, Dai Y, Song M, Zhou Y, Zhou J, Yan X, Shen Y (2021a) The decrease of intraflagellar transport impairs sensory perception and metabolism in ageing. Nature communications 12: 1789

Zhang Z, Danne N, Meddens B, Heller I, Peterman EJG (2021b) Direct imaging of intraflagellar-transport turnarounds reveals that motors detach, diffuse, and reattach to opposite-direction trains. Proceedings of the National Academy of Sciences of the United States of America 118

Zhao Q, Li J, Tong Y, Li Y, Han W, Li Z, Wang Y, Yin Y, Fang J, Jiang W et al (2026) TRiC folds the giant ciliary protein IFT172 via a non-canonical open-state mechanism. 2026.2003.2028.714460

